# Revisiting evolutionary rate-time relationships

**DOI:** 10.1101/2023.12.02.569704

**Authors:** Stephen De Lisle, Erik I. Svensson

## Abstract

Rates of molecular, phenotypic, and lineage diversification typically scale negatively with time interval of measurement, raising longstanding questions about time-dependency of evolutionary processes. These patterns and their potential meaning have recently re-entered evolutionary discussions. In this Perspective we revisit the general challenges in interpreting rate-time relationships. Much apparent temporal scaling of evolutionary rate is an inescapable outcome of plotting a ratio against its denominator, either directly or indirectly. Highly unlikely relationships between timescale and accumulated evolutionary change are required to produce anything other than negative rate-time relationships. Simulations reveal that constant rate evolutionary processes readily generate negative rate-time scaling relationships under many conditions, and that a range of rate-time scaling exponents can be generated by similar evolutionary processes. Reanalysis of six empirical datasets reveals unscaled magnitudes of evolution that are either unrelated to time and/or vary in their relationship with time, with over 99% of variation in rate-time relationships across six datasets explained by time variation alone. We further evaluated a recent hypothesis that evolutionary rate-time scaling reflects three modes of change, from micro- to macroevolutionary time scales using break-point regression, but we found no strong support for this hypothesis. Taken together, negative rate-time relationships are largely inevitable and difficult to interpret. In contrast, it is more straightforward to assess how evolutionary change accumulates with time.

## Main Text

Estimates of evolutionary rate vary with the time interval over which they are measured. A general pattern with faster evolutionary rate over short timescales, and slower rates over longer timescales has been recognized for decades (Gingerich 1983, Gingerich 1993, Hendry and Kinnison 1999, Gingerich 2001, Harmon, Pennell et al. 2021). Such time-dependency has been documented for nearly every possible measure of evolutionary rate (Rolland, Francisco Henao-Diaz et al. 2023), including rates of morphological and molecular evolution (Alba and Castresana 2005, Ho, Shapiro et al. 2007, Ho, Lanfear et al. 2011), and rates of lineage formation on phylogenies (Ricklefs 2006, Scholl and Wiens 2016) and in the fossil record (Foote 1994, Foote 2000, Henao Diaz, Harmon et al. 2019) (Figure 1). This puzzling and general phenomenon of widespread negative rate-time scaling relationships was recently highlighted as a major unsolved challenge in evolutionary biology, with potential explanation based on shared time-dependent features of evolution (Henao Diaz, Harmon et al. 2019, Harmon, Pennell et al. 2021, Rolland, Francisco Henao-Diaz et al. 2023). Recent technical and statistical explanations for negative rate-time scaling relationships move beyond simple arguments of artefact and focus on sampling biases (Emerson 2007, Louca, Francisco Henao-Diaz et al. 2022, Reijenga and Close 2025), such as a the “push of the past” (Rolland, Francisco Henao-Diaz et al. 2023), model mis-specification (Harmon, Pennell et al. 2021), and measurement error (O’Meara and Beaulieu 2024). The renewed interest in time-dependency of evolutionary rates has rekindled long-standing and controversial ideas about fundamental disparities between micro- and macroevolution (Eldredge and Gould 1972, Gould and Eldredge 1977, Charlesworth, Lande et al. 1982, Gould 1984, Gould and Eldredge 1993, Hansen and Houle 2004).

**Fig. 1.**
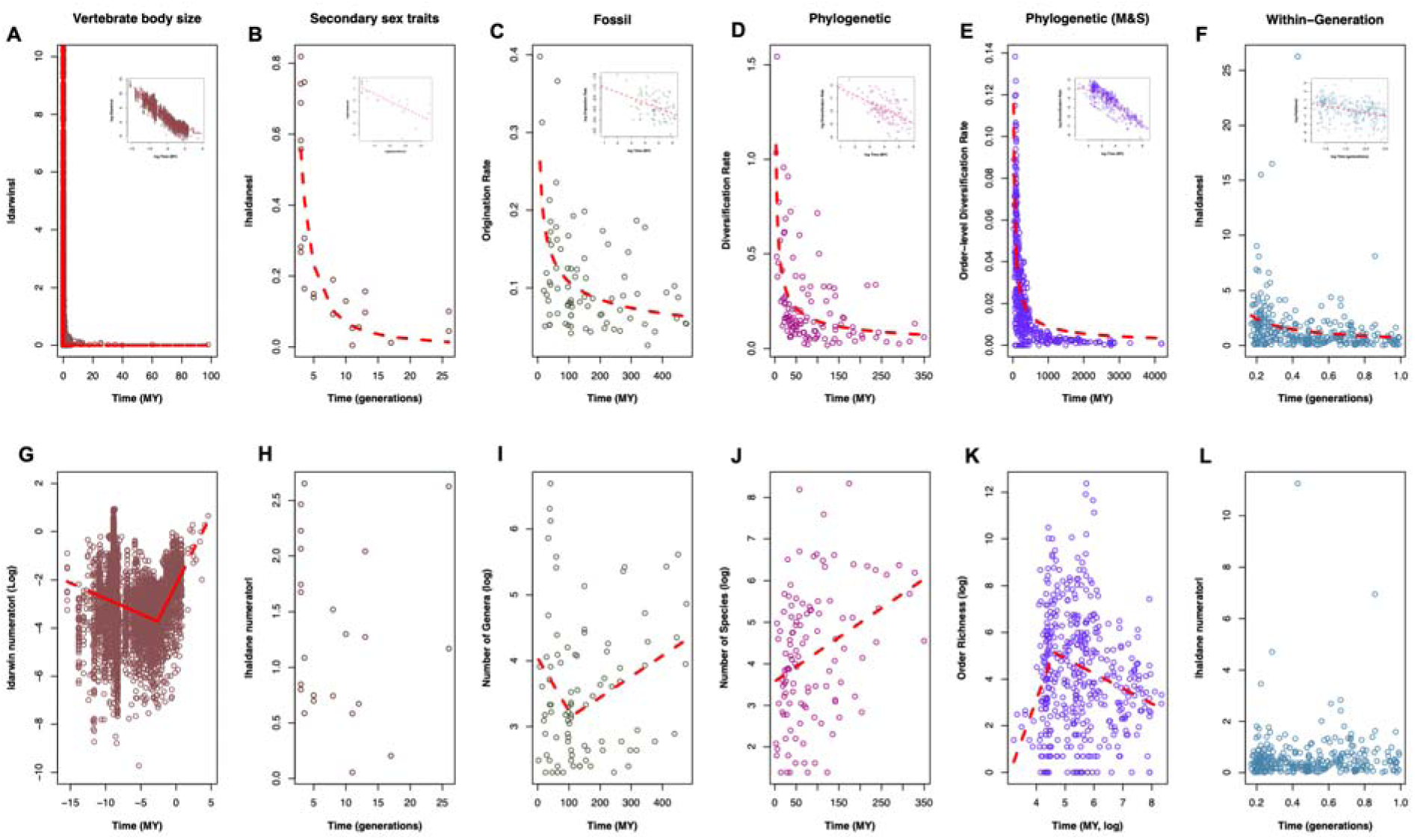
Evolutionary rate-time relationships reflect null or variable underlying relationships between evolutionary change and time. Panels A-F show evolutionary rate plotted against the timescale of measurement (Rolland, Francisco Henao-Diaz et al. 2023), with dashed red lines showing the least squares fit of the function *rate = a\**time^b^ and inset panels showing the linear relationship on a log-log scale. Panels G-L plot the amount of evolutionary change (i.e., a rate numerator) against time frame for the same datasets, with dashed red lines showing the linear breakpoint regression, if statistically significant. Panels A and G show vertebrate body size evolution (Uyeda, Hansen et al. 2011), Panels B and H show microevolution of secondary sex traits (Svensson 2019), C and I show diversification of genera from fossil lineages (plants, mammals, and marine animals)(Henao Diaz, Harmon et al. 2019), Panels D and J show diversification data estimated from time-calibrated molecular phylogenies (Henao Diaz, Harmon et al. 2019), while Panels E and K show a larger database of phylogenetic diversification rates (Scholl and Wiens 2016) where rates were inferred directly from age and richness data (Magallon and Sanderson 2001). Panels F and L show within-generation morphological changes (mainly from fish harvesting). All nonlinear negative relationships plotted in A-F were statistically significant, whereas relationships in G-L were variable, with statistically significant breakpoints found for morphology, fossil lineage accumulation, and one estimator of phylogenetic diversification, and a positive linear relationship for another estimator of phylogenetic diversification. No significant relationship was found for microevolution of sex traits (H) or the within-generation changes in panel L. Sample sizes for each column: A N = 5392 records of evolutionary change, B N = 23 records of evolutionary change, C N = 90 lineages, D N = 114 clades, E N = 434 Orders, F N = 302 records of within-generation change. Note that each of these datasets is an amalgamation of evolutionary events from many different lineages, and does not represent single anagenetic time series; thus decreasing numerator-time relationships are indeed possible.

Interpreting the causes and meaning of rate-time scaling relationships is difficult in light of the inevitable negative correlation generated by plotting a ratio against its own denominator (Pearson 1897, Atchley, Gaskins et al. 1976, Gingerich 1983, Prarie and Bird 1989, Jackson and Somers 1991, Arnold 2014). Despite widespread appreciation of this statistical issue, (e.g., see Box 1 in Gingerich 1983, Gould 1984, Harmon, Pennell et al. 2021) a revisit to the interpretative challenges of general rate-time relationships is timely and necessary in light of renewed interest in negative rate-time scaling. For example, in a high-profile review on bridging micro and macroevolution, Rolland et al (Rolland, Francisco Henao-Diaz et al. 2023) presented five negative rate-time scaling relationships as evidence of a divide between evolution over short versus deep time.

In this Perspective, we analyze a simple model of rate-time relationship that is purposefully agnostic to any specific underlying evolutionary process. We illustrate how evolutionary change, time, and their (co)variation contribute to generating nearly inevitable rate-time relationships.

We also re-analyzed six classical datasets that have been used in past work to demonstrate the universality of rate-time scaling. A central conclusion of this Perspective is that pervasive negative rate-time relationships say little on their own about the correspondence of evolutionary processes across disparate timescales.

## I. Background: Negative rate-time relationships are nearly inevitable

Evolutionary rate can be conceptualized as a function of evolutionary change scaled by time interval:

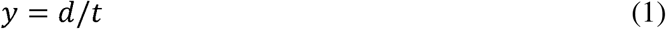

where *y* is the rate-measure, *d* is some measure of evolutionary change (e.g., change in population mean phenotype, base pair substitutions, log change in number of species in a clade) and *t* is some measure of time interval over which change was observed or inferred (e.g. generations, or years) and noting that rates are undefined at *t* = 0. Note that we use “time” and “time interval” relatively interchangeably, although it is important to realize many archetypical datasets of rate estimates contain time intervals sampled from different lineages (see below).

Some classical estimates of evolutionary rate for morphological traits are calculated directly by dividing phenotypic change with time, the latter typically measured in generations or years. For example, darwins (Haldane 1949) 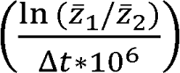, where *t* is units of years, and haldanes (Gingerich 1993) 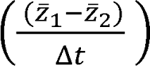, where *z* is units of pooled within-population standard deviation and *t* is units of generations, have been used to describe morphological rate, and other types of rate estimators measure per-lineage origination rate of fossil lineages (Foote 2000) and phylogenetic net diversification rate (Magallon and Sanderson 2001). Other rate estimates are obtained from more sophisticated statistical models of evolution (Nee, May et al. 1994, Nee 2001, FitzJohn 2012, Rabosky, Grundler et al. 2014). Regardless, all evolutionary rate measures are, per definition, scaled by time (e.g., waiting times between observed branching events on a phylogeny are a direct component of the likelihood in maximum likelihood models of birth-death diversification (Nee, May et al. 1994)). Thus, the conceptualization of evolutionary rate in equation 1 is general, if not precise.

The inevitable mathematical dependence between rate and time depicted in equation 1 is an obvious challenge in interpreting the time-dependency of evolutionary rate, and has been widely appreciated in evolutionary biology, both for rates of morphological and molecular evolution (Gingerich 1983, Hendry and Kinnison 1999, Roopnarine 2003, Estes and Arnold 2007, Uyeda, Hansen et al. 2011, Arnold 2014) and lineage diversification (Rabosky 2009, Rabosky 2010, Wiens 2011, Rabosky, Slater et al. 2012, Rabosky 2016, Rabosky and Benson 2021). There has also been considerable attention given to related issues outside of or tangential to evolutionary biology, including geology (Sadler 1981, Schlager, Marsal et al. 1998), paleontology (Foote 1994, Sheets and Mitchell 2001) ecology (Kenney 1982) and conservation (Wiens and Saban 2025).

When considering the role for ratio effects in contributing to our expectation for a preponderance of negative rate-time relationships, it is useful to ask: under what scenarios do we expect null or positive relationships between rate and time? Consider rate as a function of time,

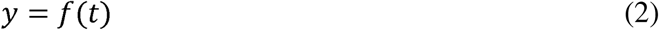

Where *y* is evolutionary rate (as in eqn 1). We also consider the amount, or magnitude, of accumulated evolution (rate numerator) as a function of time,

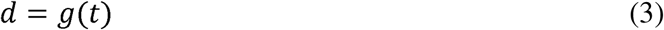

where *d* is the accumulated evolutionary change. Note that in most analyses that assess rate-time relationships, point estimates of rates from different lineages spanning different time intervals are combined in a single analysis, and in this case the function *g(t)* need not be strictly increasing even for absolute values of morphological change. Rates, *y =d/t*, will be constant over time when *d* scales linearly with *t*, i.e. *g’(t)* is a positive constant over the timescale in question, with prime denoting the first derivative. Rates will increase with time, i.e. f’(t) >0 when *g’’(t)* > 0; that is, rates increase over time when the amount of evolutionary change is an accelerating function of time. Noteworthy is that under a null relationship between *d* and *t*, such that *g’(t) = 0*, the logarithmic regression of rate versus time (log(y) ∼ μ + β*log(*t*)) will yield a negative slope estimate of β = −1 (Gingerich 1983). This can be seen most easily by expressing equation 3 as a power function, e.g., d = *t*^0^, in which case we can describe the regression of log(y) ∼ β*log(*t*) as log(*t*^0^/*t*) ∼ β*log(*t*); solving for β = log(*t*^-1^)/log(*t*) = −1. This negative slope estimate provides one cutoff of interest in assessing rate-time scaling relationships: log-log regressions of rate vs time that are shallower than or not significantly different from −1 are consistent with either a null relationship between evolutionary change and time (time independence) or an increasing relationship between accumulated change and time, respectively.

The relationships between equations 1-3 are illustrated in Figure 2, which shows the relationship between rate and time. Figure 2A illustrate differing relationships between rate numerator and time illustrated in Figure 2B for a range of power functions. There are few theoretical models that predict the type of accelerating relationship between magnitude of evolution and time that are needed to produce a positive relationship between evolutionary rate and time. Neutral and adaptive models of phenotypic and molecular evolution predict either no relationship with time or linear accumulation (constant change) with time (Morgan 1998).

**Fig. 2.**
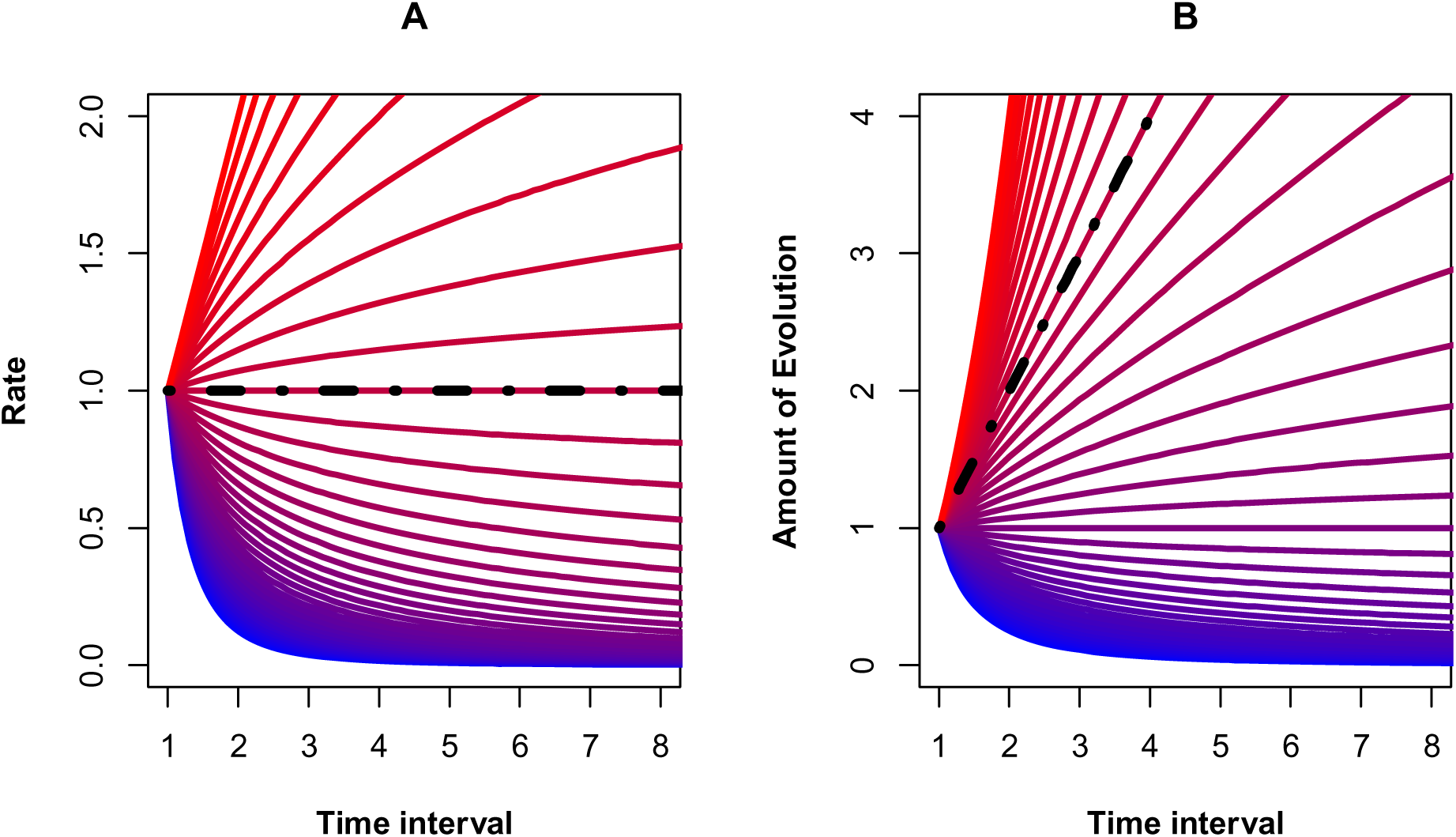
Extraordinary relationships between the accumulated amount of evolution and time are required to produce positive rate-time relationships. Panel **A** shows the relationship between rate and time for various functions of the rate numerator and time, which are plotted in the same color in Panel **B**. The black line is a reference point, where there is a null relationship between rate and time. Thus, each line in Panel **A** corresponds to a specific relationship between rate numerator and time, plotted in Panel **B**; only cases of an accelerating relationship between amount of evolution and time are expected to produce positive rate-time scaling relationships. We assume for this example a form of the relationship, *amount of evolution = time^b^*, where the function was plotted for b ranging from −2 (blue) to 2 (red). Similar general patterns are observed for other functions of change and time. Note that because we are considering these relationships in light of large amalgamations of data from disparate groups (e.g., Figure 1), as opposed to within-lineage time series, negative relationships between amount of evolution and interval time are in principle possible, even if unlikely.

Models based on adaptive landscape theory predict a slowdown in evolutionary rates for morphological traits, but a recent study of thousands and extinct and extant species did not find any empirical support for such a slowdown. i. e. no evidence for time-dependent evolutionary rates (Quintero 2025). Similarly, birth-death models of lineage accumulation also predict either a linear or a decelerating accumulation of log lineage number through time under most conditions. Even models of time-dependent lineage diversification make no prediction about accelerating log lineage number versus interval time across a sample of clades. Moreover, lineage-through-time (LLT) plots for single clades diversifying under such a process can produce complex patterns (Paradis 2015) that are consistent with several different underlying diversification processes (Louca and Pennell 2020).

Some scenarios in nature could potentially lead to accelerating change with time and thus generate positive relationships between evolutionary rates and time (Figure 2). Such scenarios include “diversity begets diversity” models of lineage diversification (Nee, May et al. 1994, Emerson and Kolm 2005), evolution by positive frequency-dependent selection (Chouteau, Arias et al. 2016), runaway sexual selection driven by female choice (Fisher 1930, Lande 1981, Kirkpatrick 1982), “tipping points” in the evolution of reproductive isolation (Nosil, Feder et al. 2017) and populations crossing critical thresholds followed by phase transitions between alternative stable states (Scheffer 2009). Tracking of a constant-rate moving optimum can generate short-term positive rate-time scaling until a population reaches its stable expected lag (Lynch and Lande 1993, Bürger and Lynch 1995) (elaborated below). How common such scenarios are in nature is an open empirical question. However, given the transient nature of self-reinforcing evolutionary processes driven by positive feedbacks (Lande 1981, Kirkpatrick 1982), we suspect that positive rate-time relationships are unlikely to be sustained over longer macroevolutionary time scales and should quickly reach stable equilibria and thus be largely transient. Rapid and self-reinforcing evolutionary processes are unlikely to proceed unchecked over millennia (Scheffer 2009), as would be required to generate positive rate-time relationships in empirical data such as that shown in Figure 1.

We can decompose the covariance between rate and time, focusing on the log-scaled case, where

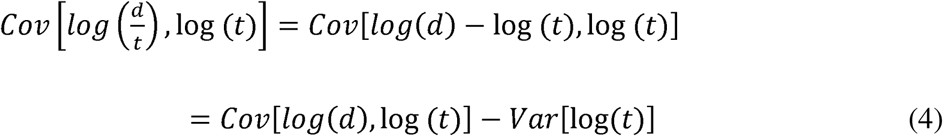

Which shows that the relationship between rate and time is governed both by the variance in time interval as well as the relationship between accumulated change (*d*) and time. Importantly, the variance partitioning in equation 4 makes it possible to evaluate the contribution of change-time relationships in explaining rate-time relationships. If rate-time relationships generally carry meaningful information about how evolutionary change accumulates with time, we expect *Cov[log(d), f=t, log (t*)] to be an important contributor to explaining variation in *Cov[log(d/t), log (t)]*. We can transform the covariance to a regression coefficient by multiplication by Var[log(t))^-1^, which results in β*_y,t_* = β*_d,t_* −1, and thus further illustrating that if there is no relationship between accumulated change d and time (such that β*_d,t_* = 0) then the expected log-log regression of rate and time, β*_y,t_*, is equal to −1.

### Simulations

Simulations of classical constant-rate evolutionary models reveal that negative rate-time associations readily emerge from these existing models. Phenotypic evolution by random genetic drift results in a Brownian motion process wherein phenotypic variance across lineages increases linearly with time (Figure 3A), and estimated evolutionary rate is time-independent when calculated for an entire clade (3E inset panel). Even this simple constant rate model results in a negative scaling relationship between lineage-specific estimates of evolutionary rate and time (Figure 3E), resulting in a scaling exponent of – ½ in a log-log regression of rate versus time (Hansen 2024).

**Fig. 3.**
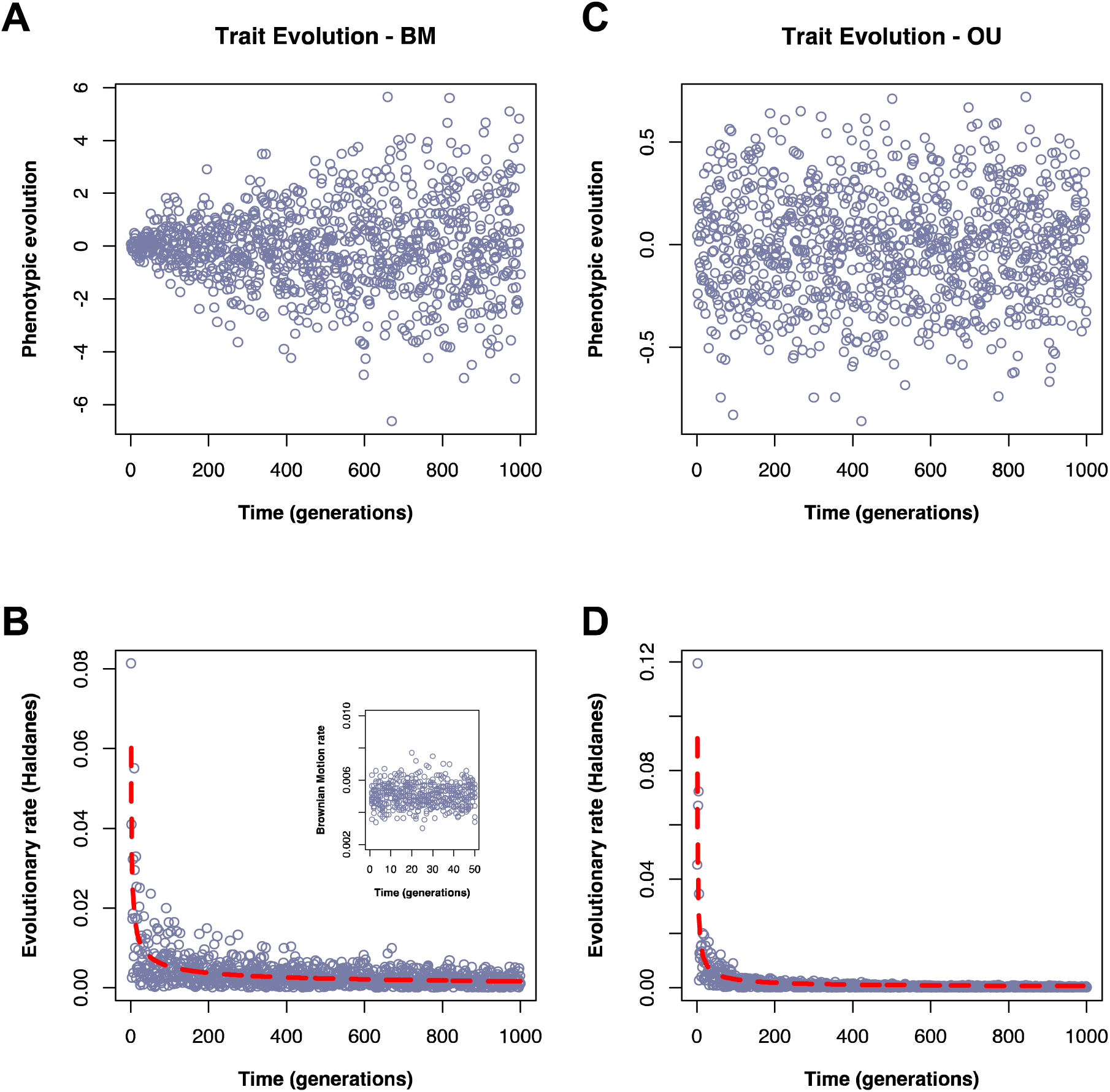
Simulations of constant-rate evolutionary models reveal persistence of negative rate-time relationships in resulting data. Panels **A** and **B** show a simulation of phenotypic evolution under random genetic drift in the absence of natural selection, which produces a Brownian motion process (Felsenstein 1988) where variance among lineages accumulates linearly with time (**A**). Although the rate of drift is constant (**B** inset), we will nevertheless obtain a negative scaling relationship between the lineage-specific per-generation rate of evolution (**B**). Panels **C** and **D** show a simulation of phenotypic evolution under drift and stabilizing natural selection, an Ornstein Uhlenbeck process (Lande 1976). This process generates no relationship between time and divergence (**C**), but again a strong negative scaling relationship with per-lineage per-generation evolutionary rate and time and hence a similar negative relationship (**D**).

Lineage specific rates are even more strongly associated with time under an adaptive model of natural selection towards a stationary optimum (an Ornstein-Uhlenbeck, or OU, process), where a null relationship between time and divergence (Figure 3B) generates a strong negative relationship between per-lineage rate and time (Figure 3F) with a log-log slope of −1 over long timescales (Hansen 2024), reminiscent of empirical data from vertebrate body size evolution (Figure 1). However, we note that a stationary OU model readily produces negative rate-time scaling exponents that deviate from −1 when sampling lineages evolving under different values of genetic variance and strength of stabilizing selection (Figure 4). Adding any movement of the optimum in an OU process produces dramatic shifts in the form of rate time scaling yet also indicates that a range of models incorporating stabilizing selection readily produce scaling relationships similar or identical to that expected under pure genetic drift (Figure 4).

**Fig 4.**
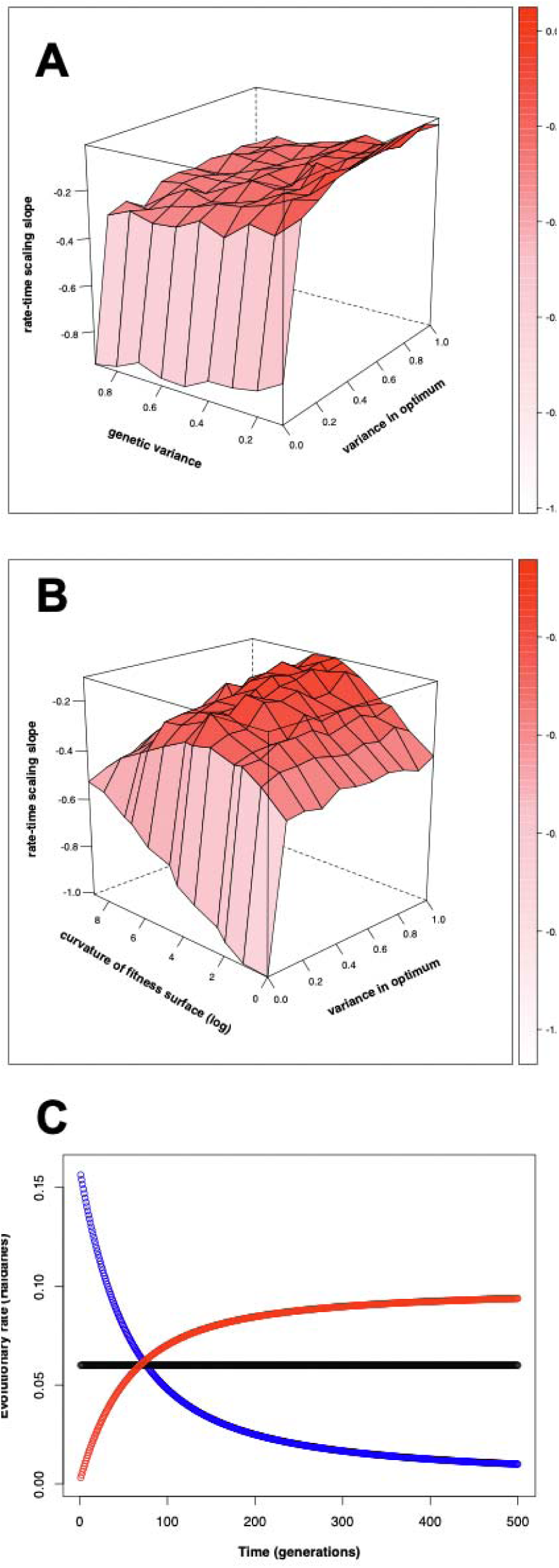
Heterogeneity in rate-time scaling exponents under different forms of morphological evolution. Panels **A** and **B** show simulated evolution under drift and stabilizing selection; an OU process. When there is zero variance in the optimum, this is a stationary OU process generating a scaling relationship of −1 over this timespan (9000 generations) only when genetic variation is relatively high (Panel **A**), and selection is strong (a low value of ω and thus a strong curvature of the fitness surface, (Panel **B**). Any variance in the optimum leads to a dramatic shift in the scaling relationship. Noteworthy is that a wide range of parameter space, corresponding to models of pure drift as well as models of pure fluctuating selection, can all produce a range of similar scaling relationships particularly in the range of 0 to −1/2. For each parameter value in **A** and **B**, the average scaling slope is shown from 10 simulation runs of 36 lineages each spanning a time range of 1-9000 generations. Panel **C** shows evolutionary rate versus time for three forms of directional evolution. In black is the rate estimate versus time for evolutionary change under the breeders equation, Δz = Gβ. Blue shows the rate estimate for evolution towards an optimum that shifted 5 phenotypic standard deviations at generation 0, assuming the same form of stabilizing selection as in Figure 4B, which is high over short timescales but decelerates as the population adapts. Red shows the relationship for evolution towards an optimum that moves at contant rate of .1 standard deviations per generation, starting at generation 0 with a population on the optimum. In this model the accelerating rate-time relationship is due to the lag between evolutionary rate and rate of peak movement that equilibrates when the population reaches its equilibrium lag behind the optimum(Lynch and Lande 1993, Bürger and Lynch 1995). All of these models produce directional evolutionary change, yet only for one specific model, Δz = Gβ, which is often thought to capture the main features of the two other models of directional change, do we see a lack of time scaling.

Well established constant-rate models of lineage accumulation can also lead to the appearance of negative associations between rate and time when sampling is incomplete, due to ratio effects. In principle Yule rates do not inherently show time dependence when estimates of zero rates, which rarely result in phylogenies of more than a single taxon, are included, yet such species-depauperate clades are rarely included in actual data, and non-zero rates show negative time-scaling even under a constant-rate pure-birth diversification process (O’Meara and Beaulieu 2024). Of course, the Yule model is not a realistic description of biological diversification, since extinction has clearly played a major role in the history of life. Yet such a combination of sampling bias and ratio effects also generate apparent negative rate-time relationships under birth-death processes, which remain to some degree even after conditioning the likelihood of the birth-death model on known sampling biases (Louca, Francisco Henao-Diaz et al. 2022).

## II. Empirical examples reveal variable patterns of accumulated change despite pervasive negative rate-time scaling

We revisited and reanalyzed six published empirical datasets of evolutionary rate estimates that have featured centrally in recent discussions of rate-time scaling relationships (Scholl and Wiens 2016, Svensson 2019, Sanderson, Beausoleil et al. 2021, Rolland, Francisco Henao-Diaz et al. 2023). We first perform regression analyses to calculate rate-time and rate numerator-time scaling relationships, the former largely recapitulating the analyses of the original studies, the latter of which is more rarely presented. We then performed the novel variance decomposition presented in equation 4, which essentially represents a metanalysis of these rate estimators.

### Vertebrate body sizes: evaluating Hansen’s (2024) “three modes” of evolution

For our first example, vertebrate body size evolution (Figure 1A, G), past work (Estes and Arnold 2007, Uyeda, Hansen et al. 2011, Arnold 2014) has shown that body size changes are largely unrelated to time, and in time-series of extant populations there was no detectable change (Sanderson, Beausoleil et al. 2021). Although a log-log regression of rate vs time yields a slope that is nominally steeper than −1 (β = −1.019, 95% CI −1.031, −1.007), which suggests no accumulation of change with interval time, there is a significant breakpoint in the relationship between morphological change and time (*F*_2, 5388_ = 207.2, *P* < 2.2 x 10^-6^, breakpoint = −2.63 log MY (0.072 MY), SE = 0.147). The result is first a weak negative relationship between time and evolutionary change, followed by an increasing slope after the breakpoint (Fig. 1A, G; or shallower negative slope on the log scale, Figure 1A inset panel), identical to that found by Uyeda et al (2011). Thus, for body size evolution, amount of change is largely unrelated to time interval except over the deepest periods of evolutionary time (Uyeda, Hansen et al. 2011). Quintero (2025) reached a similar conclusion in a recent compilation of rates of morphological evolution among thousands of extinct and extinct species and no evidence for time-dependency.

Next, we applied breakpoint regression models to the log-log relationship between body size evolutionary rate and time to test Hansen’s (Hansen 2024) recently proposed “three modes” of evolution. Hansen (2024) argued that rate-time scaling may be near null over short (microevolutionary) time intervals, approach −1 over intermediate time intervals, and relax to around −1/2 over deep macroevolutionary time intervals. To evaluate the evidence for this hypothesis we fit progressively more complex breakpoint regression models of log absolute darwins versus log time interval (Fig. 5). There was statistical support (significantly reduced AIC) for increasing complexity up to seven breakpoints (Figure 5, Table S1), with ΔAIC of over 1200 between the best fit model and one with only two breakpoints (i.e. three modes). Further, examination of the scaling slopes from a model with only two breakpoints reveals scaling slopes that all differ significantly from the values proposed by Hansen (2024); β_1_ = −0.72 (95% CI −0.78, −0.67), β_2_ = −1.19 (95% CI −1.22, −1.17), β_3_ = −0.23 (95% CI −0.32, −0.13).

**Figure 5.**
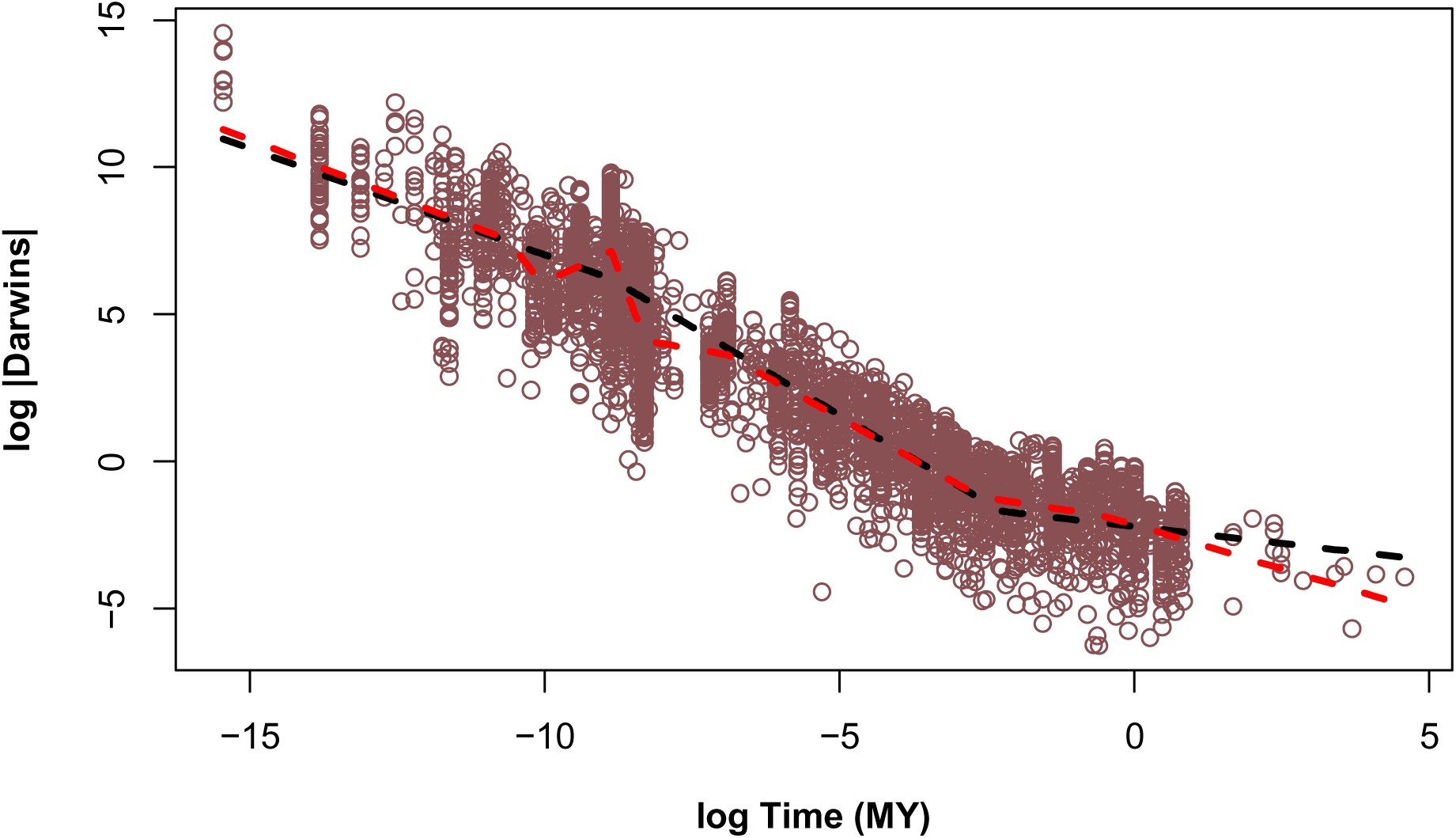
No evidence for three distinct modes of evolution revealed in vertebrate body size evolutionary rate-time scaling. Shown is the log rate-time scaling relationship for vertebrate body size data (Uyeda, Hansen et al. 2011). The black line shows the breakpoint regression with two breakpoints, corresponding to a model of three “modes” of rate-time scaling (Hansen 2024), while the red line shows the best-fit breakpoint regression (see Table S1).

### Secondary sexual traits

Svensson (2019) presents a compilation of contemporary rates of evolution of secondary sexual traits in the wild. These traits include morphology, color, and behavior traits in fish, birds, mammals, and insects that have been shown to be a target of sexual selection and span a time range of 3-26 generations. We find a negative relationship between rates and generation time interval (Figure 1B); a log-log regression of absolute haldanes versus time reveals a slope that does not differ from −1 (β = −1.306, 95% CI −1.825, −0.786), corresponding to a null relationship between haldane numerator and time (Figure 1H). This finding of a −1 scaling relationship over timespans spanning a handful of generations also contradicts the proposal for null scaling relationships over such short microevolutionary intervals (Hansen 2024).

### Fossil lineages

Similar results are seen in patterns of lineage accumulation in the fossil record (Figure 1C, I), where a strong negative rate-time relationship (found by (Henao Diaz, Harmon et al. 2019) and replotted here) reflects a weak and variable relationship between evolutionary change and time (breakpoint regression *F*_2, 86_ = 3.199, *P* = 0.046, breakpoint = −109 MY, SE = 33.13). However, this breakpoint regression model was not a significantly better fit than a model with only an intercept; *F*_3, 86_ = 2.6, *P* = 0.057. Consistent with this finding, a log-log regression between rate and time yields a slope that is significantly shallower than −1 (β = −0.24, 95%CI −0.36, −0.11) and no significant evidence for a breakpoint (*F*_2, 75_ = 1.71, *P* = 0.18), indicating accumulation of diversity with interval time.

### Phylogenetic trees

On phylogenetic trees (Figure 1D, E, J, K), there is also a consistent striking negative relationships between rate and time (as found by the original studies; Scholl and Wiens 2016, Henao Diaz, Harmon et al. 2019) that corresponds to underlying variable association between diversity and time (Rabosky, Slater et al. 2012, but see Scholl and Wiens 2016). We recover a weak positive linear relationship between diversity and time in one dataset (*F*_1, 112_ = 14.17, *P* = 0.000267; Figure 1J) corresponding to a log-log regression that is significantly shallower than −1 (β = −0.53, 95%CI −0.68, −.39), indicating some accumulation of diversity with time, and a breakpoint linear relationship in another dataset (Figure 1K; breakpoint regression *F*_2, 430_ = 12.67, *P* = 4.474 ×10^-6^, breakpoint = 4.6 log MY, SE = 0.108) corresponding to a log-log regression of rate and time with a slope that is not significantly different from −1 (β = −1.04, 95%CI −1.1, − 0.98), indicating the accumulated diversity and interval time are decoupled.

### Within-generation change

Interestingly, we even recover a nonlinear negative relationship between evolutionary rates and time when examining phenotypic change that occurred within a timeframe less than the average generation time of the study organism (here primarily long-lived fish; Figure 1F). This within-generation dataset had a log-log regression of rate and time with a slope that was not significantly different from −1 (β = −0.83, 95%CI −1.08, −0.58), indicating that there is no accumulation of change with interval time for intervals less than one generation, as we may expect since such change is likely an outcome of non-accruing environmental effects. This final example demonstrates the appearance of a rate-time scaling relationship for a case that obviously defies evolutionary explanation based on between-generation change. This conclusion is further strengthened by the fact that the slope estimate does not significantly differ from the expectation under a random relationship between rate numerator and time. That is, scaling phenotypic change by time to calculate rate generates a non-linear negative relationship with time even for this within-generation change (Figure 1F, L), that is likely to reflect phenotypic plasticity, culling, or other non-evolutionary within-generational environmental effects.

### Decomposing the covariance between rate and time across datasets

We can then decompose the covariance between rate and time for each of these datasets into its components associated with variation in time and covariance between accumulated change and time, as in equation 4. A caveat in this exercise is that not all of the rate estimates are calculated directly as in equation 1, for example phylogenetic lineage diversification rates in (Henao Diaz, Harmon et al. 2019), and so we do not expect all of the variance in rate-time relationships to be explained by these two components. Nonetheless, this analysis reveals that 99.5% (39% with morphology data dropped) of the variance in rate-time relationships can be explained by variance in time alone (*F_1,4_* = 844.5, *P* = 8.3×10^-6^, Figure 6A), while change-time covariance alone explains only 10.6 % (29% with morphology data dropped) of the variance in rate-time relationships (*F_1,4_* = 0.47, *P* = 0.53, Figure 6B). Placing both predictors in a multiple regression reveals support for variance in time (*t* = −28.4, *P* = 9.5×10^-5^, df = 3), but not covariance between change and time (*t* = 1.13, *P* = 0.34, df = 3), as a significant predictor of rate-time relationships. While all covariance relationships will depend strongly on variance in their constituent variables, this analysis reveals the noteworthy contrast in lack of meaningful contribution of covariance between accumulated change and time.

**Fig. 6.**
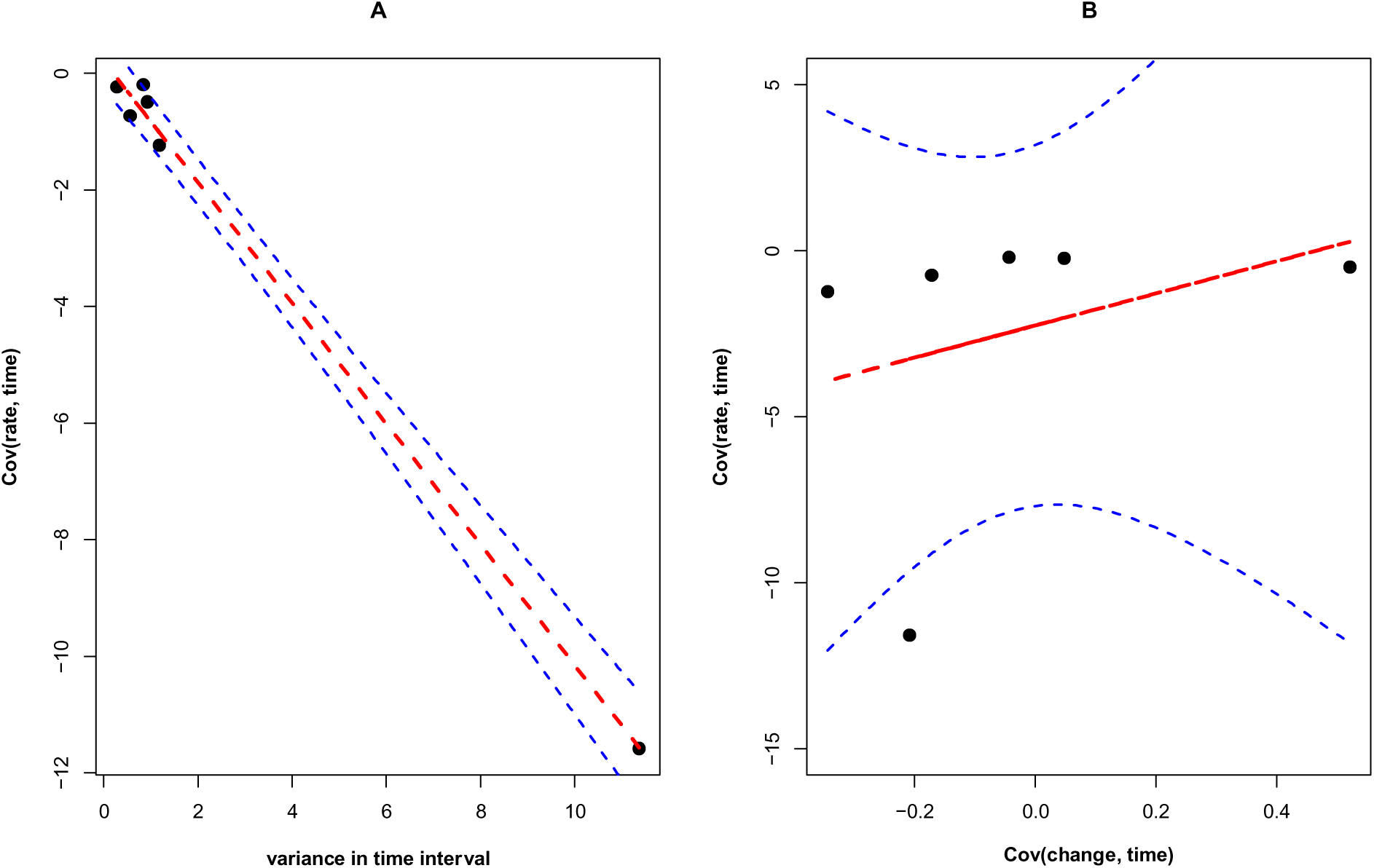
Time interval determines rate-time relationships across datasets. The covariance between rate and time in the six empirical datasets shown in Figure 1 is almost entirely explained by variance in time (Panel **A**). Covariance between change and time, also expected to be a component of the covariance between rate and time, is not significantly associated (Panel **B**). Lines and 95% confidence intervals are from linear models fit separately to each; a multiple regression with both var(time) and cov(change, time) as predictors yielded equivalent conclusions (see text).

## Discussion

In this perspective we have explored the problem of why all estimated evolutionary rates appear to scale negatively with time, and what information may be gleaned from these rate-time relationships. First, it is readily apparent there will only exist a limited range of relationships between the amount of evolution and time that generate anything other than negative rate-time relationships. Second, revisiting several independent datasets of evolutionary rates illustrates that existing, widely documented negative rate-time scaling relationships clearly suggest weak and inconsistent relationships between the amount of evolution and timescale of measurement, with no evidence of consistent time-dependency in evolutionary change across datasets. Our novel analysis of these datasets reveals that variation in rate-time scaling relationships are primarily driven by variation in timespans rather than patterns of accumulated evolutionary change. Given that most common process-based models of constant-rate evolutionary change can readily generate a range of negative rate-time scaling relationships, the ubiquitous negative rate-time relationships in empirical data is hardly surprising. Although some of these points have been discussed before, our revisit to the problem unequivocally illustrates the challenge in interpreting negative rate-time scaling relationships, timely in light of their re-emergence in recent literature (Henao Diaz, Harmon et al. 2019, Harmon, Pennell et al. 2021, Rolland, Francisco Henao-Diaz et al. 2023).

### How meaningful are rate-time scaling relationships?

Despite clear interpretive challenges inherent in evolutionary rate-time relationships, evolutionary processes must of course play a driving role in generating them (Kinnison and Hendry 2001, Ho, Lanfear et al. 2011, Henao Diaz, Harmon et al. 2019, Harmon, Pennell et al. 2021, Louca, Francisco Henao-Diaz et al. 2022, Rolland, Francisco Henao-Diaz et al. 2023), and so there is justified interest in understanding the drivers of rate-time scaling relationships beyond the inevitable qualitative expectation of a negative relationship (Harmon, Pennell et al. 2021).

Analyses of rate-time relationships can perhaps provide new insights when coupled with explicit null hypothesis tests against scaling parameters expected under various evolutionary process models (as can be done for the case of morphological evolution; Hansen 2024, Holstad, Voje et al. 2024).

Here, we performed such a test and found no strong evidence for the three modes of morphological evolutionary rate-time scaling proposed by Hansen (Hansen 2024) neither in our reanalysis of an existing database of vertebrate body size evolutionary rate (Figure 5) nor in our finding of a −1 relationship between microevolutionary rate and time (Figure 1b). Moreover, it is challenging to construct meaningful null expectations for other types of rate-time scaling patterns, such as lineage diversification. For example, in their assessment of negative scaling of diversification rates, Henao Diaz et al. (2019) constructed a null expectation by simulating birth-death trees for the same age ranges as their empirical set of trees, under the mean values of birth and death rate observed for the oldest groups in the study. While such an approach is useful in identifying cases of negative rate-time scaling beyond what may be an inevitable outcome from our imperfect statistical models (but see Reijenga and Close 2025), for many types of rate estimate we lack quantitative rate scaling predictions. One point of our paper is that, without some robust quantitative expectation for a rate scaling relationship, the existence of a negative relationship between empirical rates and time is not very informative.

Nonetheless, it will often be difficult to reconcile quantitative rate-time scaling patterns with specific evolutionary process models, especially in large datasets of evolutionary rate estimates spanning disparate lineages. For example, although a stationary OU model can generate a −1 scaling relationship of rate with time (Hansen 2024), but this is only strictly true over moderate timescales when selection is strong and genetic variance is moderate to high.

Similarly, evolution by Brownian motion produces a scaling relationship of −1/2, but so can a random walk of the optimum under stabilizing selection, a process which itself is not strictly Brownian (in terms of trait evolution) unless all phenotypic variance has an additive genetic basis. Thus, while dramatic shifts in rate-time scaling relationships do imply shifts in the mode of evolution (Hansen 2024), many similar rate-time scaling relationships may be generated by many disparate processes. Similarly, widespread decoupling of clade size and clade age has led to proposals that rate-time relationships and measures of net diversification rate carry little conceptual value (Rabosky 2009, Rabosky 2010, Rabosky, Slater et al. 2012, Rabosky and Hurlburt 2015, Rabosky and Benson 2021), although counterpoints have been made (Wiens 2011, Harmon and Harrison 2015, Kozak and Wiens 2016). Similar arguments could be made for morphological evolution: if evolutionary change rarely accumulates in a consistent manner with time, what is the value of scaling change by time in the first place?

### Parameter vs summary rates: A distinction without difference?

In general, we should be surprised when estimated evolutionary rates *do not* show negative time scaling. However, some estimators of evolutionary rate do not show the types of inevitable time scaling relationships we have discussed; for example, clade-level estimates of Brownian motion rate appear to be less affected. Moreover, constancy of evolutionary rates, and thus linear accumulation of change with time is sometimes observed at the molecular level (Morgan 1998, Bromham and Penny 2003). Indeed, in their review, Harmon et al. (see also Hunt 2012, 2021) draw a distinction between “summary rates” and “parameter rates”; the former being descriptive summary statistics, such as haldanes, while the latter are inferences of instantaneous rate parameters estimated from a process-based model of evolutionary change, with Brownian motion rate or birth-death lineage diversification rate being two well known examples. In this distinction, and barring inadequate models or data, it has been proposed that parameter rates from constant rate models should be free from inevitable time-scaling (Hunt 2012, Harmon, Pennell et al. 2021). This need not be strictly true, since an estimator of a population parameter can be the maximum likelihood estimator yet still be biased in certain contexts (a classic example is Pearson’s correlation coefficient, which is the MLE of the correlation for a bivariate normal population, but demonstrates small-sample bias; Fisher 1915). Many maximum likelihood estimators of parameter rates are ultimately simple functions of amount of evolution and time, and noteworthy is that even most summary rates could in fact be considered parameter rates under appropriate evolutionary models (Harmon, Pennell et al. 2021). The haldane, for example is a parameter rate under one very simple model of directional trait change (constant directional selection) but shows positive and negative time scaling in different process-based manifestations of directional evolution models (see Fig. 4C). Given that we can never know the true process model producing an evolutionary dataset all our models will always be wrong to some degree (Box 1976). It remains to be seen if there is a general distinction between these categories of rate estimate in regard to prevalence or interpretability of time scaling.

### Analysis of rate numerators

Relationships between the magnitude of evolutionary change (e.g., phenotypic change or the log number of new lineages arising) and time can be assessed directly, as we and others have done (Hendry and Kinnison 1999, Uyeda, Hansen et al. 2011, Svensson 2019). Breakpoint regression (Schlager, Marsal et al. 1998, Hendry and Kinnison 1999, Uyeda, Hansen et al. 2011, Clapham and Karr 2012) can be used to identify significant breakpoints in patterns of accumulated evolutionary change across time. Such an approach avoids the problem of statistical confounders between rate and time interval, at least when applied to analysis of change-time relationships, and also allow for the possibility of identifying shifts in the form of rate-time scaling relationships (Hansen 2024). Although analysis of rate-time and change-time plots present the same total information, uninformed comparisons of rate-time relationships will readily generate the appearance of shared features of the evolutionary process. Our analysis revealed that variation in evolutionary rate-time relationships across datasets revealed little information about variation in how evolutionary change accumulated with time.

## Conclusions

A negative rate-time relationship is nearly inevitable, and the meaning and importance of negative rate-time relationships for the correspondence of microevolution and macroevolution is still unclear. We end with a final example that illustrates this point. Consider the widely-appreciated fact that microevolutionary quantitative genetic models of random genetic drift are inconsistent with the amount of evolutionary change in deep time (Lynch 1990, Hansen and Houle 2004, Houle, Bolstad et al. 2017). We often find far less evolutionary change in deep time than expected via drift alone, based on patterns of standing extant genetic variance. Yet even if phenotypic divergence in deep time is consistent with drift under observed levels of standing variance, we would still expect to see a negative scaling relationship between lineage specific rates of evolution and time interval (Figure 3). This fact alone is underappreciated and highlights that great caution is needed in interpreting negative rate-time relationships.

## Methods Details

### Data

We analyze existing data on rates of vertebrate body size evolution (Uyeda, Hansen et al. 2011), which consists of data on log-fold change in body size as well as time interval over which this change occurred. For phylogenetic lineage diversification, we analyze two datasets, one in which diversification rates were estimated from the branching pattern of species-level phylogenies (Henao Diaz, Harmon et al. 2019), and another larger dataset (Scholl and Wiens 2016) where diversification rates were estimated from clade diversity and age information. For fossil data, we analyze an existing dataset (Henao Diaz, Harmon et al. 2019) of emergence rates of new genera. For each dataset, we plot the standard measure of rate against the timeframe over which that rate was measured, fitting exponential linear models by nonlinear least squares (Rolland, Francisco Henao-Diaz et al. 2023). We also plot a measure of the rate numerator against time for each of these datasets, and fit breakpoint linear regressions to identify significant linear relationships between amount of evolution and time, as well as any potential shifts in the relationship between evolution and time. For each dataset, we fit nonlinear models of the form *rate = a*time^b^* using nonlinear least squares (nls()) in R, as well as breakpoint regression using the segmented package. We compared models fits of an ordinary linear regression to a model with a single breakpoint, and also when appropriate to a null model containing only an intercept, using the anova() function in R. Data presented in Figure 1 were obtained from (Henao Diaz, Harmon et al. 2019) (available at https://github.com/fhenaod/evol_time_scaling/tree/main/data), (Uyeda, Hansen et al. 2011)(Dryad doi:10.5061/dryad.7d580), (Scholl and Wiens 2016), and (Sanderson, Beausoleil et al. 2021) (database available at: https://proceeddatabase.weebly.com/).

### Simulation

We simulated three different constant rate evolutionary processes. The aim of our simulations was not to evaluate the performance of statistical models used to estimate rates. Rather, our aim was simple to examine scaling patterns observed in the simplest cases of simulated constant rate evolution. For morphological evolution, we simulated random genetic drift for 1-1000 generations assuming a heritability of 0.50 and a population size of 100. We also simulated evolution under genetic drift and stabilizing selection, with the same demographic and genetic assumptions but assuming weak selection (*ω*^2^ = 15) towards an optimum. Each simulation was started with the population at its optimum. For morphological evolution, we plotted rates in units of Haldanes (Gingerich 1993), which was natural as we simulated evolution on the scale of phenotypic standard deviations. We then repeated our simulations of OU evolution but exploring a range of parameter values for genetic variance, the strength of selection (curvature of the fitness surface) and the stability of the optimum. For the later, we sampled changes in the optimum from a normal distribution centered at zero and exploring different values of the variance, producing a random walk of the optimum.

**Table S1.**
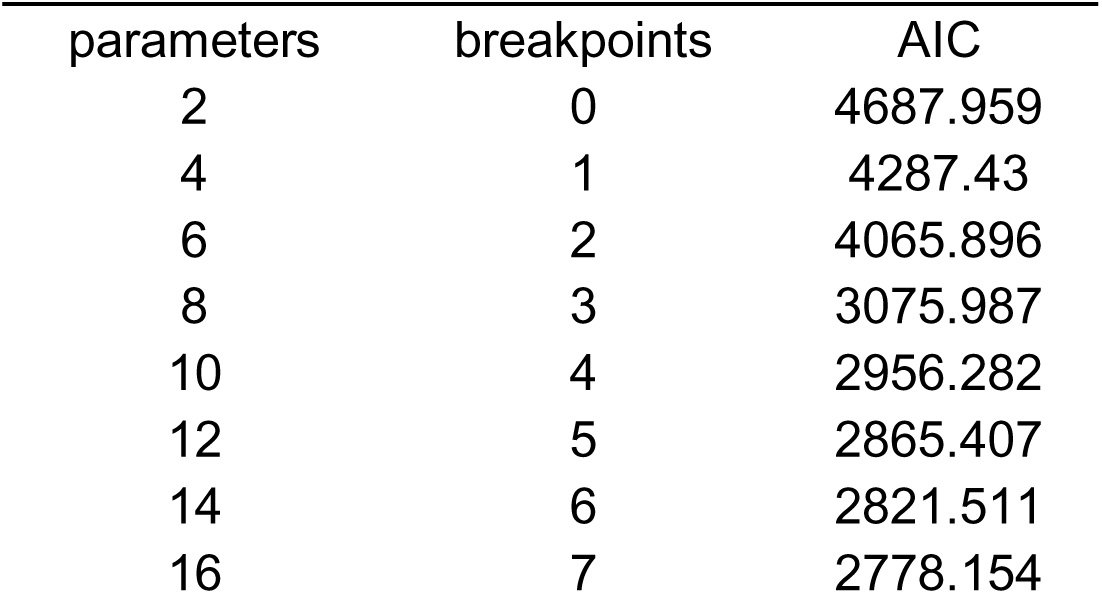
Model fits for testing Hansen’s “three modes” hypothesis. . Note that models

| parameters | breakpoints | AIC |
| --- | --- | --- |
| 2 | 0 | 4687.959 |
| 4 | 1 | 4287.43 |
| 6 | 2 | 4065.896 |
| 8 | 3 | 3075.987 |
| 10 | 4 | 2956.282 |
| 12 | 5 | 2865.407 |
| 14 | 6 | 2821.511 |
| 16 | 7 | 2778.154 |

## Notes

### Competing Interest Statement

The authors have declared no competing interest.

### Summary of Updates

This has been revised to contain updated analyses, including addition of new data, removal of redundant text, and substantial revision to the text. The main conclusions remain unchanged.

